# A temperature dependent trophic cascade modifies temperature dependence of ecosystem function

**DOI:** 10.1101/233916

**Authors:** Jessica Garzke, Stephanie J. Connor, Ulrich Sommer, Mary I. O’Connor

## Abstract

Ecological communities and their ecosystem functions are sensitive to temperature, and aquatic habitats worldwide continue to experience unprecedented warming. Understanding ecological effects of warming requires linking empirical evidence to theories that allow projection to unobserved conditions. Metabolic scaling theory and its tests suggest that warming accelerates ecosystem functions (e.g., oxygen flux), yet this prediction apparently contradicts community-level studies suggesting warming is a stressor that can reduce ecosystem function. We sought to reconcile these predictions with an experimental test of the hypothesis that cascading trophic interactions modify the temperature-dependence of community structure and ecosystem fluxes. In a series of independent freshwater ecosystems exposed to a thermal gradient, we found that warmer temperatures strengthened the trophic cascade increased and indirectly changed community structure by altering grazer species composition and phytoplankton biomass. Temperature-driven community shifts only modestly affected the temperature dependence of net ecosystem oxygen fluxes. Over the 10 °C thermal gradient, NPP and ER increased ∼2.7-fold among ecosystems, while standing phytoplankton biomass declined by 85-95%. The exponential increase in oxygen flux over the thermal gradient, as well as monotonic declines in phytoplankton standing stock, suggested no threshold effects of warming across systems. We also observed temperature variation over time, within ecosystems. For phytoplankton biomass, temporal variation had the opposite effect to spatial variation, suggesting that within-community temporal change in community structure was not predicted by space-for-time substitution. We conclude that food chain length can modify effects of temperature on ecosystem fluxes, but that temperature can still have continuous and positive effects on ecosystem fluxes, consistent with patterns based on large-scale, macroecological comparisons. Changes in community structure, including temperature dependent trophic cascades, may be compatible with prevailing and predictable effects of temperature on ecosystem functions related to fundamental effects of temperature on metabolism.

**Statement of authorship:** JG & MIO designed the study, MIO & US provided materials, JG & SJC performed research and collected data, JG performed zooplankton analysis, SJC performed phytoplankton analysis, JG & MIO performed modeling work, analyzed data output, and wrote the first draft, and all authors contributed substantially to reviews

## Introduction

Understanding how warming associated with climate change affects species interactions and communities is one of the most pressing current ecological challenges. Two leading conceptual frameworks, the Metabolic Scaling Theory (MST) and community ecology, produce very different predictions for community-scale responses to environmental temperature. Metabolic scaling theory predicts that the relative temperature dependence of major metabolic functions explains a large fraction of variation in ecosystem fluxes and biomass stocks (López-Urrutia *et al.*, 2006; Enquist *et al.*, 2007; Wohlers *et al.*, 2009; Yvon-Durocher *et al.*, 2010; O’Connor *et al.*, 2011; Yvon-Durocher *et al.*, 2012; Barneche *et al.*, 2014). This signal of temperature dependence of fundamental metabolic rates emerges over broad climate gradients and comparisons among independent ecosystems (López-Urrutia *et al.*, 2006; Anderson-Teixeira *et al.*, 2008; Yvon-Durocher *et al.*, 2012) and suggests that energetic constraints of two highly conserved metabolic processes (oxygenic photosynthesis and aerobic respiration) may drive responses to environmental temperature change at the ecosystem, community and population levels (Gillooly *et al.*, 2001; Yvon-Durocher *et al.*, 2010; 2012; Padfield *et al.*, 2016).

The metabolic scaling approach to understanding community level responses to temperature change is challenged by an apparent incongruence between effects predicted by the temperature-dependence of metabolic functions such as photosynthesis and respiration, on the one hand, and the often large effects of temperature on species interactions, on the other. Temperature affects species interactions via population and bioenergetic dynamics, often producing non-intuitive patterns in species’ abundances (Beisner & Peres-Neto, 2009; O’Connor, 2009; Kordas *et al.*, 2011; Dell *et al.*, 2013; Gilbert *et al.*, 2014). Numerous warming experiments have shown that warming alters the abundance of interacting consumers and resources and the strength of top down control of community structure (Hansson *et al.*, 2012; Shurin *et al.*, 2012). It has even been suggested that MST is not relevant to understanding community change in response to warming (Tilman *et al.*, 2004; Brauer *et al.*, 2009). Whether demographic or bioenergetic changes at the community scale are constrained by highly conserved metabolic thermal asymmetries, or represent additional and potentially confounding effects of temperature on species interactions, remains unresolved. This lack of resolution implies that we have no accepted framework for determining which patterns observed in simple mesocosm experiments can be extrapolated to projections for effects of climate change in nature.

If the effects of temperature on consumer-resource interactions are constrained by the temperature dependence of photosynthesis and respiration (Allen *et al.*, 2005), net ecosystem effects of warming may be largely independent of demography or species interactions. In other words, though temperature-dependent trophic interactions can alter density, biomass and species composition of consumers and primary producers (Beisner *et al.*, 1996; Petchey *et al.*, 1999; O’Connor *et al.*, 2009; Yvon-Durocher *et al.*, 2010; DeLong *et al.*, 2015), there is little evidence that these community changes cause temperature-dependence of ecosystem-level oxygen fluxes to deviate from expectations based on the temperature dependence of photosynthesis and respiration at macroecological scales (López-Urrutia *et al.*, 2006; Anderson-Teixeira *et al.*, 2008; Yvon-Durocher *et al.*, 2010; 2012). The best evidence to support the importance of community shifts in modifying ecosystem level responses to temperature comes from experimental tests of gross primary production of periphyton (in the absence of grazing) across a thermal gradient in streams (Padfield *et al.*, 2017). These results showed that species composition shifts can reduce the effect of temperature at the ecosystem level, compensating for effects of temperature on total productivity. However, we do not yet have evidence that trophic interactions can play such a role, although local-scale experiments with a few species suggest that the strength or presence of strong top-down control by consumers can influence ecosystem level energy flux (Schindler *et al.*, 1997); one likely pathway, therefore, through which species interactions could modify the effect of temperature on net ecosystem fluxes is if trophic cascades increase in strength at higher temperatures. The strength of trophic cascades depends on primary production and the magnitude of herbivore density and/or behavioral responses to predation (Polis *et al.*, 2000; Schmitz *et al.*, 2003), in addition to the activities and density of predators themselves. Temperature dependent trophic cascades could therefore disrupt relationships between temperature and ecosystem fluxes mediated by direct effects of temperature on per capita metabolic rates.

Here, we tested whether temperature-dependent top-down control on herbivore and algal abundance and composition can alter the effect of temperature on ecosystem function (oxygen flux, phytoplankton standing stock). We experimentally quantified change across a broad temperature gradient (10 °C) and compared ecosystem and community states across this gradient (photosynthesis and respiration). The purpose of the broad temperature gradient, which exceeds forecasted warming over the coming century, is to test the functional response of warming at the community and ecosystem scales to allow comparison with theoretical predictions about this relationship. This question gets at the heart of the larger question of whether energy and material fluxes in ecosystems can be adequately predicted at the local scale in terms of temperature constraints on fundamental metabolic rates (photosynthesis, respiration), or whether community dynamics render such metabolic-theory-derived predictions insufficient to the point of not useful (Brown *et al.*, 2004; Tilman *et al.*, 2004; Brauer *et al.*, 2009).

### MST framework and hypotheses

We express our hypotheses in terms of testable relationships between temperature and ecosystem function. One common approach to understanding how temperature affects communities and ecosystems is to ‘scale up’ from individual physiological processes and species-specific traits. This approach requires a high burden of information about each species’ thermal traits, and this need for detailed species-level information prohibits such scaling-up for most ecosystems. The macro-ecological framework of metabolic scaling theory provides an alternative approach, in which relatively little information (in this case, temperature dependence of the fundamental and highly conserved metabolic processes photosynthesis and respiration) is applied to whole systems with many individuals and species to understand the aggregate functional response to temperature change. In this framework, whole-organism metabolic rates (e.g., oxygen flux) and related biological functions for organism *i* have been described as following a power-law dependence on body mass and exponential (Boltzmann-Arrhenius) dependence on temperature (Gillooly *et al.*, 2001; Brown *et al.*, 2004):

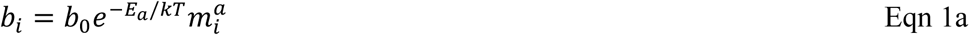

in which activation energy (*E*_a_, in eV) captures temperature (T, in Kelvin) effects on per capita metabolic response (*b_i_*) for individual *i, k* is the Boltzmann constant (eV/K), *b_0_* is a normalization constant independent of body size and temperature, *m_i_* corresponds to the body mass of an individual *i*, and *a* is the allometric scaling factor. This model is a special case of a more complex equation that allows each species to follow a thermal performance curve, often described by a modified version of the Sharpe-Schofield equation, in which performance declines at high temperatures. However, for multi-species systems, the exponential model performs well (Padfield *et al.*, 2017). This exponential model has been extended to produce a first-order expectation for the effects of temperature on ecosystem-level rates (*B_R_*):

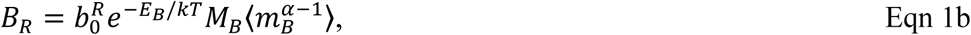

in which – *Eb* is the ecosystem-level temperature dependence term for ecosystem rate *R*, *M*B** is total biomass of the community, and 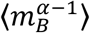 is a weighted average biomass. Together, 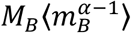 represents ‘mass-corrected’ biomass, which is a measure of the total metabolic capacity of the biomass pool in the ecosystem (Yvon-Durocher & Allen, 2012). Here, we tested the effects of temperature on independent (isolated) experimental ecosystems with distinct thermal histories, sharing a common initial species pool. Following Eqn 1b, our hypotheses about how the mean temperatures of these independent ecosystems lead to patterns in ecosystem function across systems can be stated as: for communities closed to immigration and ecosystem closed to resource inputs other than light:

i. The temperature dependence (–*E_B_* = –*E_NPP_*) of net primary production (*B_NPP_ = B_NPP_*) is predicted by the temperature dependence of photosynthesis (*E_PS_* = *E_NPP_* = −0.32 eV) for autotroph-dominated communities,
ii. Across systems, the temperature dependence –*E_ER_* of net ecosystem respiration *B_ER_* is predicted by the temperature dependence of respiration *E_ER_* =*E_R_* = -*0.65* eV)
iii. These temperature dependence coefficients do not vary with food chain length (Fig. 1).

These hypotheses that net ecosystem fluxes vary with temperature with the activation energies associated with per capita metabolic rates (photosynthesis and temperature) assume that minimal changes occur in total mass-corrected autotrophic and heterotrophic metabolic biomass over the thermal gradient within each trophic treatment (Yvon-Durocher *et al.*, 2010; Yvon-Durocher & Allen, 2012; Yvon-Durocher *et al.*, 2012). Past experiments on species interactions suggest this assumption is not supported, so, following community ecological theory and empirical evidence, we consider two alternate hypotheses:

*(iv)* total phytoplankton biomass standing stock (*M_B_*) declines with temperature and this effect varies with food chain length reflecting a temperature-dependent trophic cascade.
*(v)* food chain length and temperature change the relative abundance of species and trophic groups within each community, but these community changes do not affect net ecosystem fluxes.

**Figure 1:**
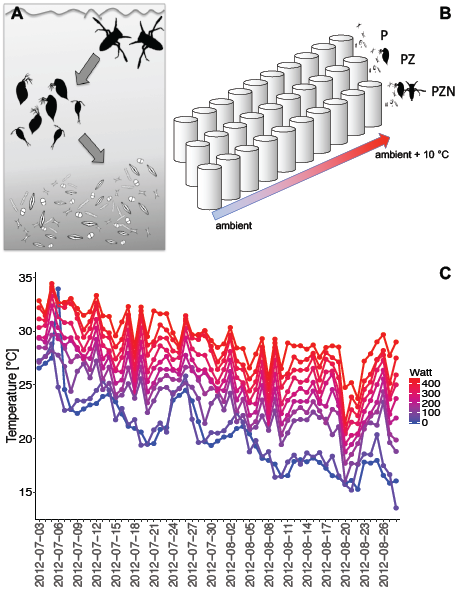
**A)** Experimental food web composition and **B**) experimental design for three trophic level systems. Food chains contained Notonectid predators (N), zooplankton (Z) grazers *Eurytemora* sp. and *Daphnia* sp., and phytoplankton (P). **C**) Ecosystem temperature and food chain length were manipulated in a regression experimental design with 10 independent systems spanning a 10 °C temperature gradient. The temperature gradient was repeated for each food chain length: 1-, 2- and 3-trophic levels. The experiment occurred over one to many generations of grazers and phytoplankton, but within one (adult) life stage of notonectid predators.

## Methods

### Experimental Design

We tested our hypotheses by manipulating temperature and food chain length in 30 independent aquatic ecosystems (Fig. 1A-B). For each food chain length (algae-only, algae-grazer or algae-grazer-predator), we maintained ecosystems at distinct temperatures in a regression design with mean temperatures ranging from ∼19 °C to ∼31 °C (Fig. 1C). The regression design allowed us to estimate slopes (*E_A_*, Eqn 1b) of response variables along a continuous temperature gradient (Cottingham *et al.*, 2005; Gotelli & Ellison, 2013) for different trophic structures by log-transforming equation 1b and fitting linear models to the continuous temperature gradient. The regression design was the right choice to compare activation energies over a broad range of the x-variable (temperature); an important test of thermal functional responses that is not possible with designs with only 2 or even three temperature levels. The three-trophic-level treatment included predators that were not a dynamic part of the system – they did not reproduce and their several month life span did not allow for demographic responses. Therefore, inferences about trophic structure are restricted to systems with dynamics in the primary producers and primary consumers, with fixed predation-related mortality imposed by a third trophic level.

We designed an experiment to test these hypotheses by comparing replicate communities across a thermal gradient. Each ecosystem experienced the same conditions (weather, seasonal variation), and differed in the average temperature of the ecosystem (Fig. 1C). Our experiment was not designed to track and measure community dynamics over time – we did not sample frequently enough for a robust test of temporal dynamics. Therefore, we aim to test our hypotheses and draw inferences based on a comparison of 30 independent ecosystems along the thermal gradient, rather than by detailed analysis of their temporal trajectories. Considerations of how ecosystems responded to temperature variation over time in this experiment would be confounded by temporal changes in community structure and successional dynamics. We focus here on how the structure and function of ecosystems varies with average temperature. Our approach thus uses a controlled experiment to mirror the application of temperature scaling models to whole ecosystem change over broad spatial scales in comparative studies (López-Urrutia *et al.*, 2006; Anderson-Teixeira *et al.*, 2008; Yvon-Durocher *et al.*, 2012).

### Experimental Food Webs

We assembled freshwater food webs in 30 outdoor mesocosms (370 L tanks) at the University of British Columbia, Vancouver, Canada (49°14’52” N, 132°13’57” W). From June 26^th^ to August 28^th^ 2012, we experimentally manipulated temperature (10 levels) and food chain length (3 levels: algae-only, algae + zooplankton, and algae + zooplankton + predator food chains, Fig. 1A-B). Initially (experiment day 7) mesocosms were inoculated with pondwater (1L) containing living algae, collected and filtered through a 64-μm sieve to remove zooplankton and larvae. Three days later, we collected zooplankton at Trout Lake, Vancouver, B.C. (49°15’23” N, 123°03’44” W), with a vertical tow net (64-μm mesh). Zooplankton were mixed in buckets to homogenize species composition, acclimated overnight to mesocosm temperatures and dead organisms removed. Initial experimental communities consisted of 25 phytoplankton taxa (S1) and predominantly 2 zooplankton taxa (the cladoceran *Daphnia sp.*, and calanoid copepod *Eurytemora* sp.), though a few cyclopoid copepods were included. To ensure colonization of grazing zooplankton, two individuals of *Daphnia* sp. and ten *Eurytemora* sp. were added to each consumer treatment (all 2- and 3-TL ecosystems). We introduced 2 individual notonectid predators *(Notonecta undulata)* on July 3^rd^, 2012 (experiment day 7) to 10 3-TL tanks. Notonectids generate trophic cascades by suppressing zooplankton (Mcardle & Lawton, 1979). Notonectids did not reproduce during the experiment, and we replaced dead notonectids during the experiment with similar-sized individuals from the same source population.

### Abiotic and biotic conditions

Mesocosms were filled with municipal water and left for one week to allow chlorine to evaporate before organisms were introduced (Kratina *et al.*, 2012). We added 160-μg NaNO_3_ L^-1^ and 10-μg KH_2_PO_4_ L^-1^ to each tank (16:1 N:P) on experiment day 7. Water was heated with submersible aquarium heaters (50, 100, 150, 200, 250, 300, 350, 400, 450 Watt) to increase temperature above ambient daily temperature. Temperature differences among tanks were consistent throughout the course of the experiment (Fig. 1C). Mesocosms were covered with two layers of window screen to minimize colonization by other invertebrates. Water levels were maintained by natural precipitation and weekly additions to maintain volume. The spatially randomized assignment of temperature and trophic treatments eliminated systematic variation in negligible allochthonous carbon inputs.

### Plankton Sampling and Analysis

Weekly, we sampled phytoplankton, chlorophyll *a*, zooplankton, and oxygen concentrations. We sampled algal assemblages in 100-mL water samples collected from ∼40-cm below the surface. We counted and identified cells using the Utermöhl sedimentation method (Utermöhl, 1958) and estimated chlorophyll *a* concentration using a Trilogy fluorometer (Turner Designs). Phytoplankton were identified and counted to species or taxon level by inverted microscopy. We collected depth-integrated zooplankton samples (10 L water filtered through a 64-μm); the filtered water returned to mesocosms. Plankton was fixed with Lugol’s iodine solution (5%). Under 10x magnification, we counted and identified zooplankton to genus, measured standard length for all development stages in week 8. We measured oxygen concentrations using YSI-85 oxygen sensor (Yellow Springs Instruments, Yellow Springs, Ohio, USA).

### Estimation of biomass and fluxes

We estimated carbon biomass of phytoplankton (*M_P_*) by converting chlorophyll *a* concentrations using 0.05 μg chlorophyll *a* / μg C at 295K, and a temperature dependence of this ratio of −0.001 °C^-1^ (Geider *et al.*, 1997). For zooplankton (grazers and notonectids), we used length-weight ratios to convert length to carbon (S2). We did not estimate microbial or periphyton biomass.

We estimated whole ecosystem oxygen fluxes using the dissolved oxygen (DO) change technique (Marzolf *et al.*, 1994), correcting for temperature effects on oxygen saturation state. The change in DO (ΔDO) was measured over 24 hours (dawn, dusk and the following dawn) according to forecasted sunrise and sunset on measurement days (Hanson *et al.*, 2003). We assumed sunrise represented minimum daily O_2_ concentration, after which all subsequent values were greater (Yvon-Durocher *et al.*, 2010; Kratina *et al.*, 2012; Yvon-Durocher & Allen, 2012), and we assumed that sunset coincided with maximum O_2_ concentration (DO) after which all subsequent values were lower. Physical oxygen flux (mg m^-2^ d^-1^) with the atmosphere was calculated as follows: O_2flux_ = exp(O_2water_ – O_2sat_) * ln(T + 45.93) - e, where O_2water_ is the O_2_ concentration of water, O_2sat_ is the concentration the water would have if it were at equilibrium with the atmosphere (390-μatm), T is the temperature correction for O_2_ saturation, e is the pressure coefficient for elevation of study area (here 0.03) (Atwood *et al.*, 2015). We estimated NPP and ER, in μmol O_2_ L^-1^ d^-1^, as follows:

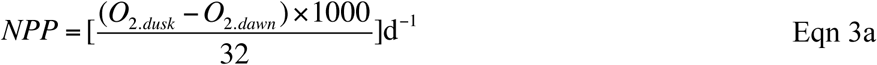

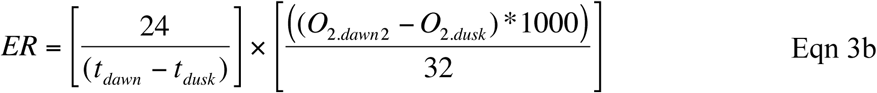

in which O_2_ is dissolved oxygen (mg L^-1^), 32 μg O_2_/μmol O_2_, and measurement time *t_i_*.

We processed and analysed the full set of response variables for weeks 4 – 9, except for zooplankton size, excluding transient bloom conditions (weeks 1-3) that were not the focus of this study.

### Statistical Analysis

We used linear mixed effects models to describe relationships between ecosystem and community response variables (Y) and temperature, a continuous predictor in our experimental design. While temperature varied over time in the experimental ecosystems, so did other conditions including ecological succession and weather. Our hypotheses are about how different average temperature conditions affect ecosystem functions, given normal environmental variation within ecosystems. Therefore, we modelled among-system responses to temperature, while also modelling within-system variation associated with temperature, and other conditions, over time. To distinguish within-tank variation from among-tank effects of temperature, we used a within-subject mean centering approach that decomposes the environmental effects into those associated with the average environment experienced over the experimental duration (‘between-tank’ effect), versus deviations of the environment in a given temperature treatment (‘within-tank temperature’ effect) (van de Pol & Wright, 2009).

We first tested a random effects model, in which the response variable (Y) for each ecosystem *j* in week *i* was modelled as a continuous response to variation in inverted ecosystem temperature (1/kT_ij_) and trophic level (TL_j_)

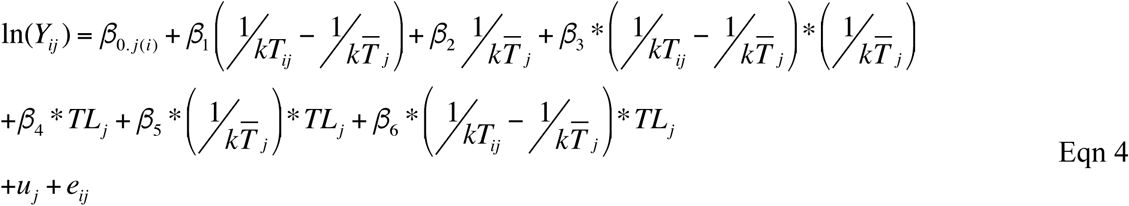

where *β_0,j(i))_* represents an intercept allowed to vary randomly among tanks. The between-ecosystem effect of temperature (*β_2_*) is estimated as the slope of ln(Y*_ij_*) on the mean value of inverse temperature for ecosystem load 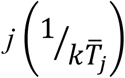 over all weeks. The within-subject (*β_1_*) effect of temperature variation over time is estimated as the slope of ln(Y*_ij_*) *vs* the deviation of the mean temperature each week 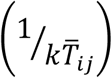 relative to 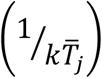. We also tested for an interaction between within-treatment temporal variation in temperature and the experimental temperature treatment (*β_3_*). To test our hypothesis that trophic structure modifies the effect of temperature on ecosystem function, we included the terms (*β_6_*) and (*β_5_*). We tested for effects of trophic structure on ecosystem function, independent of temperature, with the term (*β_4_*). Response variables were ln-transformed prior to analyses to achieve normal distributions and to linearize temperature effects for analysis and comparison with MST predictions (Eqn 1b). When modelling we centered temperature treatment (1/kTj) on the grand mean temperature *T̅* (not shown in Eqn 4) to reduce correlations between slope and intercept terms. For each response variable, we used ANOVA to determine the need for the random effect of tank by first comparing the full model (Eqn 4) with the same model without the random effect.

To test our hypothesis that food chain length (1, 2 or 3 trophic levels) modifies the estimated temperature dependence (*E_a_*, Eqn 1b) of oxygen flux and biomass, we first identified the best model for each response variable. We compared models with and without trophic level terms (*β_4_*) and interactions between *TL* and temperature (*β_5_*), and *β_6_*). We also tested models without each of the temperature terms. In total, after testing for the random effects structure, the model set included 9 models (Table 1). We ranked models using Akaike’s Information Criterion (AIC_c_) weights, and compared models using likelihood ratio tests (LRT). When two or models were considered comparable or equivalent δAIC < 2) we reported all models meeting this criterion and report averaged coefficients. We estimated activation energy and intercepts for among tank responses to temperatures by first rearranging Eqn 4 to group coefficients by temperature term (Eqn 4a):

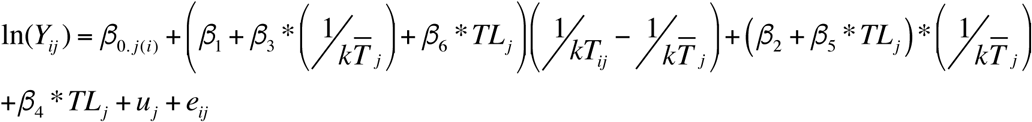

**Table 1:** Candidate models. The full model, as written in Eqn 4 in the main text, is reproduced here for convenience. The table indicates with an ‘x’ which terms were included in each candidate model.

|  | Model parameter |  |  |  |  |  |  |  |
| --- | --- | --- | --- | --- | --- | --- | --- | --- |
| Mod. | $\beta_{0,j(i)}$ | $\beta_1$ | $\beta_2$ | $\beta_3$ | $\beta_4$ | $\beta_5$ | $\beta_6$ | $\mu_j$ |
| Full.r | x | x | x | x | x | x | x | x |
| Full | x | x | x | x | x | x | x |  |
| 8 | x | x | x |  | x | x | x |  |
| 7 | x | x | x |  | x | x |  |  |
| 6 | x | x |  |  | x |  | x |  |
| 5 | x | x |  |  | x |  |  |  |
| 4 | x | x | x | x |  |  |  |  |
| 3 | x | x | x |  |  |  |  |  |
| 2 | x | x |  |  |  |  |  |  |
| 1 | x |  |  |  | x |  |  |  |
| 0 | x |  |  |  |  |  |  |  |

We estimated confidence intervals for composite terms following (Figueiras *et al.*, 1998). To analyse the effects of time and temperature on the phytoplankton community composition, we performed a Non-metric multidimensional scaling (NMDS). It is a rank-based ordination technique which is robust towards data sets where abundances are highly diverse among taxa. While NMDS does not allow assessing the effects of environmental gradients, it enables the detection of temporal patterns in the data. We used the metaMDS function in R to measure the Bray-Curtis community dissimilarities. Species abundances were square root transformed. We used R statistical software (R v. 1.0.44 R Developmental Core Team 2006) with packages MuMIn, nlme, plyr, tidyverse, broom, reshape2, lubridate, hms and zoo.

## Results

### Temperature dependence of top-down control (hypotheses iv - v)

We first present results for hypotheses iv-v, because they demonstrate top-down control that was temperature dependent in our ecosystems. Total ecosystem oxygen fluxes varied with food chain length, after controlling for temperature, reflecting top down control of ecosystem functions, as indicated by significant variation in intercept terms and the *β_4_* term in the highest ranking models for NPP and ER (Fig. 2, Table 2). Net oxygen fluxes in systems with predators tended to be more similar to systems without consumers (Fig. 2A-F), indicating that strong grazer effects on oxygen fluxes were minimized in the presence of predators by top down control.

**Figure 2:**
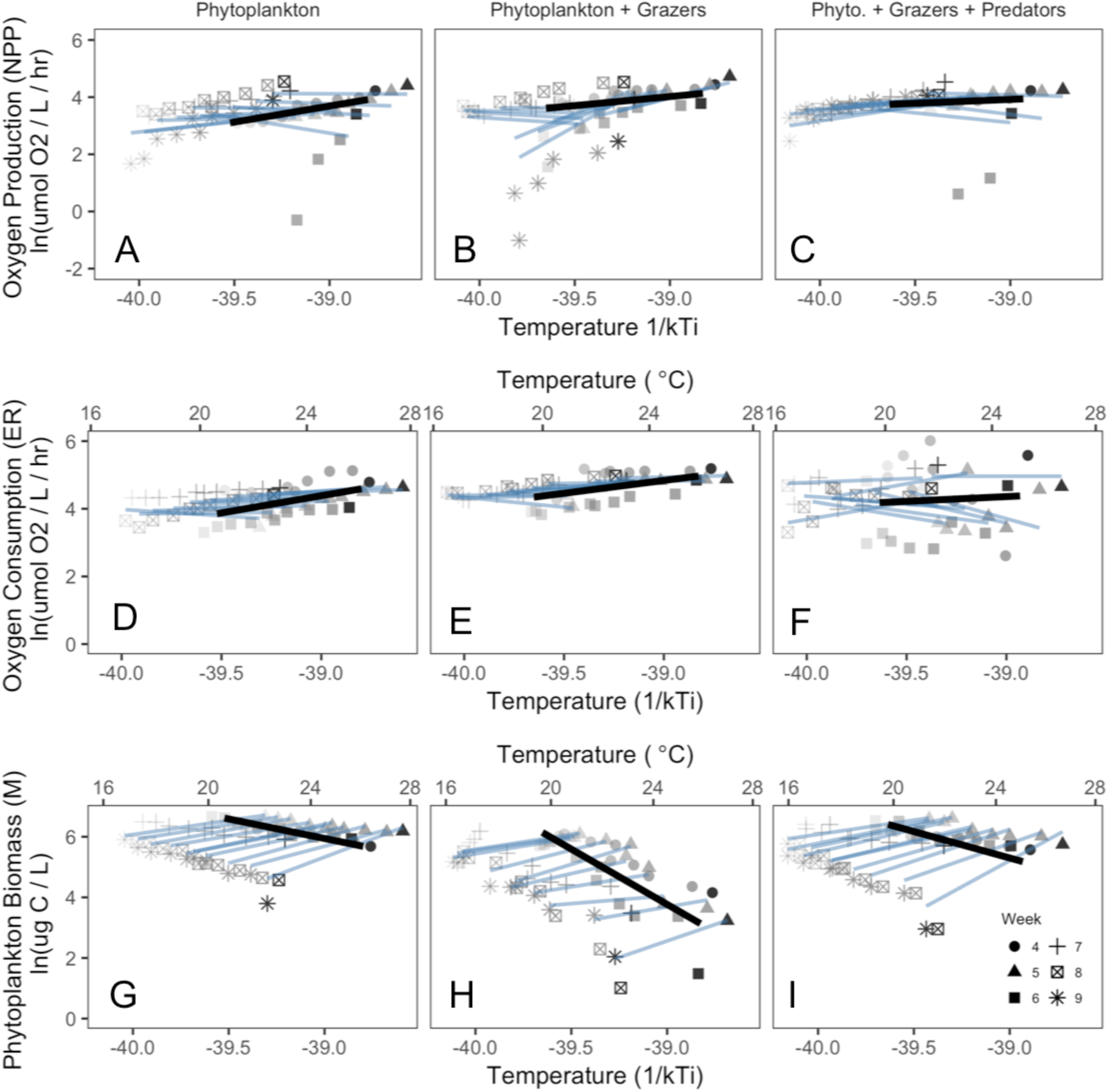
Net ecosystem oxygen flux, phytoplankton standing stock, and estimated activation energies varied over the experimental thermal gradient. **A-C**) Net primary production (NPP), **D-F**) net ecosystem respiration (ER) and **G-I**) phytoplankton standing stocks (*M_B_*) were estimated once per week (for 6 weeks post bloom) in each replicate ecosystem (n = 30). For each ecosystem (shade of grey), 6 points are shown, one point for each week (symbols). Bold lines indicate hierarchical regressions fit to among-group variation in temperature, after taking into account within-group variation temperature effects (light lines)(Table 2). Temperatures within tanks declined over time (Fig. 1C). Activation energies and confidence intervals given in Table 3.

**Table 2:**
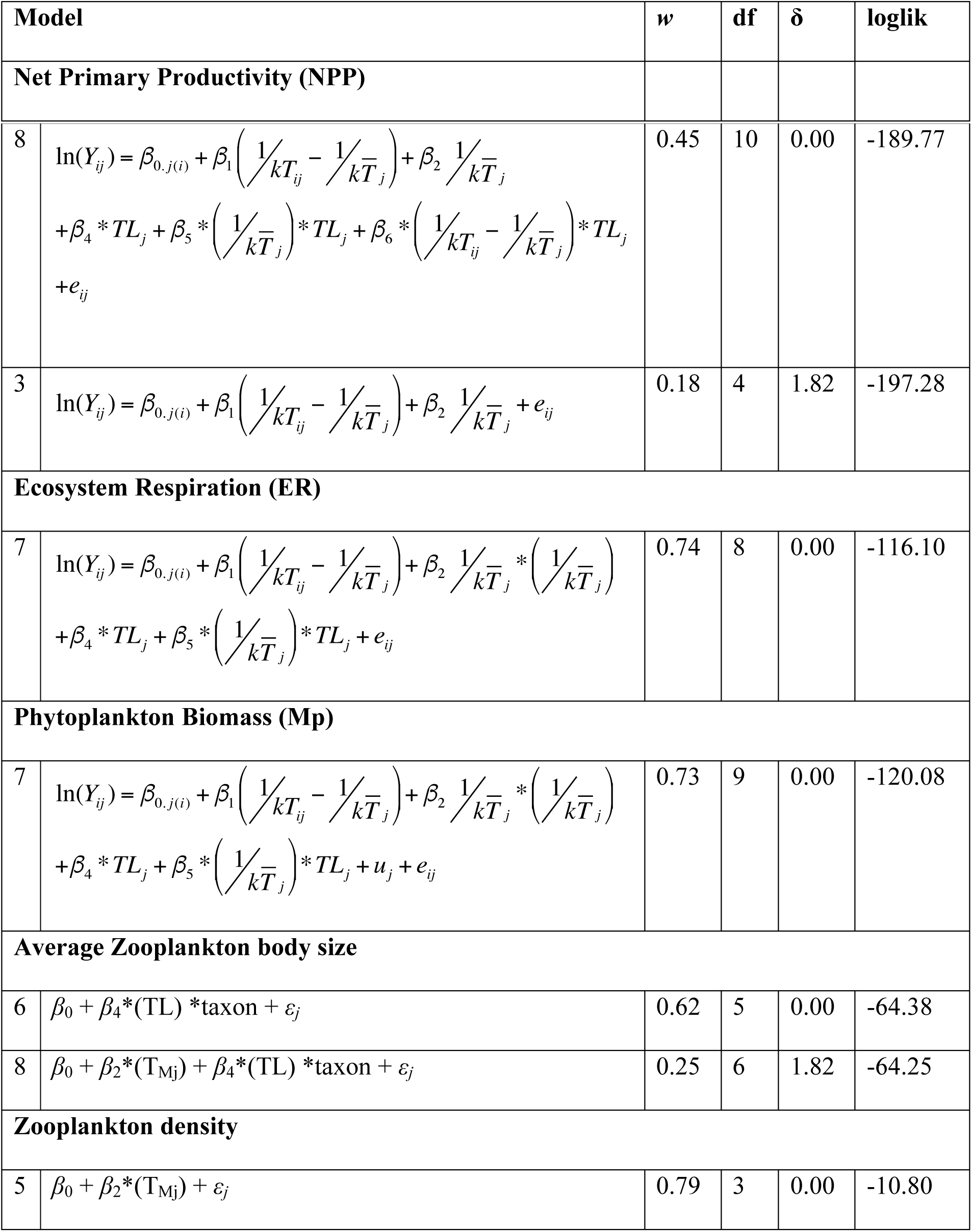
Model comparisons for effects of temperature and food chain length on biological responses based on AIC weight (*w*) and δAIC values. Response variables are modelled as in Equation 4, as functions of temperature T_ij_ for each tank *j* on week *i* relative to the mean temperature T_Mj_ for tank *j* over all weeks (T in Kelvin), and food chain length (TL). Only models ranking in the top set δAIC < 2) are shown.

| Model | $w$ | df | $\delta$ | loglik |
| --- | --- | --- | --- | --- |
| Net Primary Productivity (NPP) |  |  |  |  |

|  |  |  |  |  |  |
| --- | --- | --- | --- | --- | --- |
| 8 | $\ln(Y_{ij}) = \beta_{0.j(i)} + \beta_1 \left( \frac{1}{kT_{ij}} - \frac{1}{k\bar{T}_j} \right) + \beta_2 \frac{1}{k\bar{T}_j}$ $+ \beta_4 * TL_j + \beta_5 * \left( \frac{1}{k\bar{T}_j} \right) * TL_j + \beta_6 * \left( \frac{1}{kT_{ij}} - \frac{1}{k\bar{T}_j} \right) * TL_j$ $+ e_{ij}$ | 0.45 | 10 | 0.00 | -189.77 |
| 3 | $\ln(Y_{ij}) = \beta_{0.j(i)} + \beta_1 \left( \frac{1}{kT_{ij}} - \frac{1}{k\bar{T}_j} \right) + \beta_2 \frac{1}{k\bar{T}_j} + e_{ij}$ | 0.18 | 4 | 1.82 | -197.28 |
| **Ecosystem Respiration (ER)** |  |  |  |  |  |
| 7 | $\ln(Y_{ij}) = \beta_{0.j(i)} + \beta_1 \left( \frac{1}{kT_{ij}} - \frac{1}{k\bar{T}_j} \right) + \beta_2 \frac{1}{k\bar{T}_j} * \left( \frac{1}{k\bar{T}_j} \right)$ $+ \beta_4 * TL_j + \beta_5 * \left( \frac{1}{k\bar{T}_j} \right) * TL_j + e_{ij}$ | 0.74 | 8 | 0.00 | -116.10 |
| **Phytoplankton Biomass (Mp)** |  |  |  |  |  |
| 7 | $\ln(Y_{ij}) = \beta_{0.j(i)} + \beta_1 \left( \frac{1}{kT_{ij}} - \frac{1}{k\bar{T}_j} \right) + \beta_2 \frac{1}{k\bar{T}_j} * \left( \frac{1}{k\bar{T}_j} \right)$ $+ \beta_4 * TL_j + \beta_5 * \left( \frac{1}{k\bar{T}_j} \right) * TL_j + u_j + e_{ij}$ | 0.73 | 9 | 0.00 | -120.08 |
| **Average Zooplankton body size** |  |  |  |  |  |
| 6 | $\beta_0 + \beta_4 * (TL) * \text{taxon} + \varepsilon_j$ | 0.62 | 5 | 0.00 | -64.38 |
| 8 | $\beta_0 + \beta_2 * (T_{Mj}) + \beta_4 * (TL) * \text{taxon} + \varepsilon_j$ | 0.25 | 6 | 1.82 | -64.25 |
| **Zooplankton density** |  |  |  |  |  |
| 5 | $\beta_0 + \beta_2 * (T_{Mj}) + \varepsilon_j$ | 0.79 | 3 | 0.00 | -10.80 |

Predator-grazer-algae food chains had higher phytoplankton biomass (*M_P_*) (Fig. 2G-I) than systems with grazers but no predators, similar to algae-only food chains. This classic trophic cascade became stronger as temperature increased (Fig. 2G-I, Table 1). By week 8, predators had shifted composition of zooplankton to less effective grazers (from *Daphnia* sp. to copepods, Fig. 3B, S3), though there was no clear effect of predators on zooplankton density in week 8. The proportion of inedible phytoplankton taxa (cyanobacteria) increased over time at higher temperatures, but this change was unaffected by food chain length (Fig. 4).

**Figure 3:**
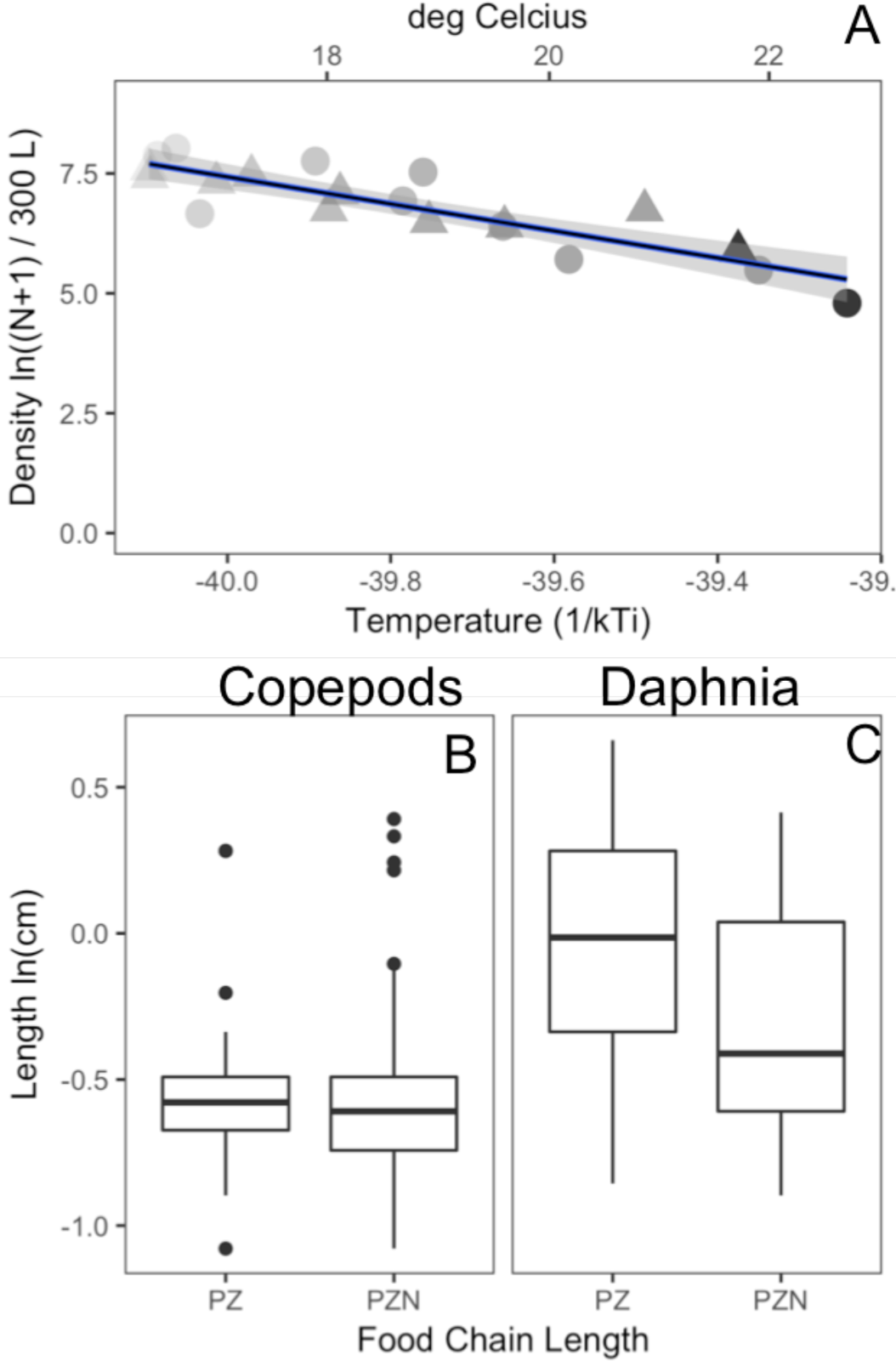
In the 8^th^ week of the experiment, zooplankton density declined with increasing temperature (A) but did not differ between ecosystems with (triangles) and without (circles) notonectid predators (Table 2). Daphnia size declined in the presence of predators, but copepod size did not, and there was no effect of temperature on body size in week 8 (Table 2).

**Figure 4.**
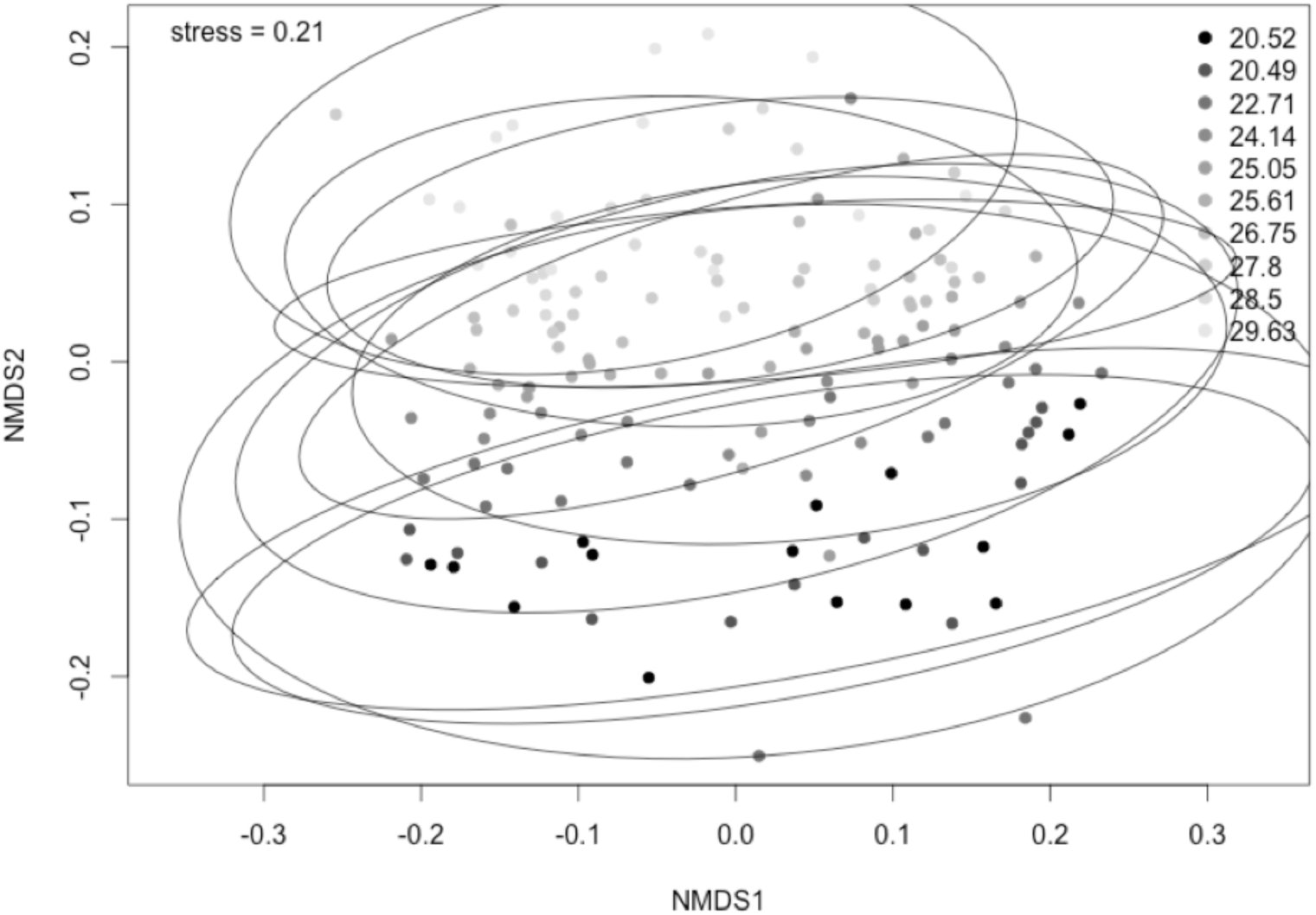
Temperature and food chain length shifted algal community composition in experimental ecosystems. Each point represents one ecosystem observed at one time, and lighter shades are communities at higher temperatures as indicated by the inset legend with mean temperature for each tank over the experimental period. Tanks sharing the same temperature treatment are indicated by an ellipse. Mesocosm phytoplankton species diversity mostly affected by time are the ones closest to the axes of the graph in the NMDS plot.

Phytoplankton biomass (*M_P_*) declined with increasing temperature (Fig. 2G-I, Table 2). Across-system effects of temperature were steepest in 2-TL systems with zooplankton grazers and no predators, with a 30-fold decline as temperature increased across systems (Fig. 2, Table 3). The trophic cascade strengthened with temperature, and we therefore do not reject hypothesis *iv.* For ecosystem-level phytoplankton biomass ln(*M_P_*), a model with random effect for tank (Full.R) performed better than the model without random effects (Full) (Full vs Full.R: p = < 0.001, S4), and subsequent analyses were done with random effects models. Within-ecosystem trends in biomass associated with temperature differed starkly from effects of temperature among ecosystems (Fig. 2). Over time, higher temperatures were associated with higher phytoplankton standing stocks within systems, though interpretation of the temperature effect is confounded by the decline in temperature over time likely associated with other temporal changes.

**Table 3.**
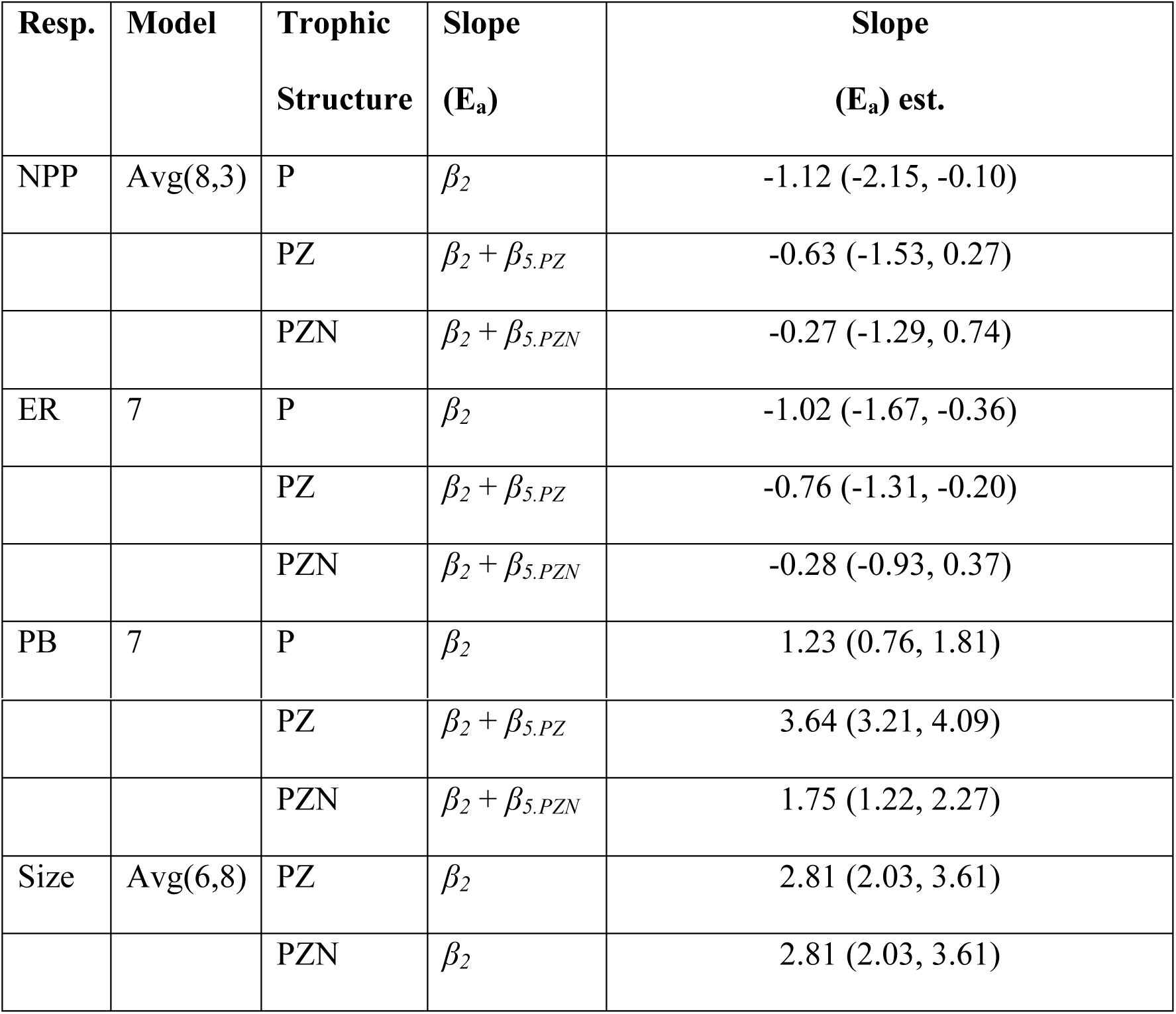
Estimates and formulae for activation energy terms.

| Resp. | Model | Trophic Structure | Slope ($E_a$) | Slope ($E_a$) est. |
| --- | --- | --- | --- | --- |
| NPP | Avg(8,3) | P | $\beta_2$ | -1.12 (-2.15, -0.10) |
| | | PZ | $\beta_2 + \beta_{5,PZ}$ | -0.63 (-1.53, 0.27) |
| | | PZN | $\beta_2 + \beta_{5,PZN}$ | -0.27 (-1.29, 0.74) |
| ER | 7 | P | $\beta_2$ | -1.02 (-1.67, -0.36) |
| | | PZ | $\beta_2 + \beta_{5,PZ}$ | -0.76 (-1.31, -0.20) |
| | | PZN | $\beta_2 + \beta_{5,PZN}$ | -0.28 (-0.93, 0.37) |
| PB | 7 | P | $\beta_2$ | 1.23 (0.76, 1.81) |
| | | PZ | $\beta_2 + \beta_{5,PZ}$ | 3.64 (3.21, 4.09) |
| | | PZN | $\beta_2 + \beta_{5,PZN}$ | 1.75 (1.22, 2.27) |
| Size | Avg(6,8) | PZ | $\beta_2$ | 2.81 (2.03, 3.61) |
| | | PZN | $\beta_2$ | 2.81 (2.03, 3.61) |

### *Trophic structure influenced temperature dependent ecosystem function* (hypotheses *i - iii)*

Our results suggest that trophic structure alters the effect of temperature on ecosystem-level oxygen flux. Across the experimental temperature gradient, net ecosystem oxygen production (NPP) increased with temperature (bold lines, Fig. 3A-C), and varied with trophic structure (Fig. 2A-C, Tables 2-3). Mean temperature dependence of NPP across ecosystems was not statistically different than the expected *E_NPP_* = −0.32 eV (consistent with hypothesis i), though the confidence intervals are wide enough to include 0 for ecosystems with consumers, and also the activation energy of respiration (-0.65 eV) (Table 3). In the absence of zooplankton the estimated across-system temperature dependence was stronger than expected: *E_NPP_* = −1.12 eV (± 1.02) (Table 3). For ecosystem-level ln(NPP), a model without a random effect for tank (Full) performed just as well as a model with the random effect (Full.R) (Full vs Full.R: p = 0.999, S5), and subsequent analyses were done with fixed effects models. The best model included an interaction between trophic level and average temperature (Table 2), suggesting that trophic structure alters across-system temperature dependence of NPP. However, the difference in the coefficients of temperature dependence among trophic treatments is not well resolved, due to the broad confidence intervals (Table 3).

Trophic structure also altered the across-system temperature dependence of ER, weakening the effect of temperature when food chains were longer. Thus we reject hypothesis *ν*. Net ecosystem respiration (ER) increased with temperature across ecosystems, consistent with the predicted *E_R_* = −0.65 eV (consistent with hypothesis *ii*, Fig. 2D-F, Table 2), and with confidence intervals that do not include the proposed temperature dependence of NPP for short and long food chain systems (Fig. 2, Table 2). For tank-level ln(ER), a model without a random effect for tank (Full) performed just as well as a model with the random effect (Full.R) (Full vs Full.R: p = 0.859, S5), and subsequent analyses were done with fixed effects models.

Within ecosystems, effects of temperature variation over time on oxygen fluxes were weaker than or comparable to among-system effects (Fig. 2), and did not depend on temperature treatment or trophic level (Table 2).

## Discussion

The growing literature of experimental tests of how warming affects interacting species aims to reduce uncertainty in projected ecological changes associated with climate change. Warming experiments have shown a wide variety of consequences for species interactions, from shifts in community composition, strengthening top-down control, and shifts in body size. However, how these responses to warming, in short-term experiments designed to test specific hypothesis, can be best related to projections for climate change impacts over broad scales of space, time and complexity is not always clear. One way to facilitate such projections is for experiments to test functional responses of ecological processes along thermal gradients, as we have done here, and to test for sensitivity of functional responses to ecological conditions. Empirically estimated functional responses that can be incorporated into theoretical models can support projections of change based on system dynamics, rather than direct extrapolation from experimental conditions, and are likely to prove the most useful in understanding ecological change with climate change (Cuddington *et al.*, 2013).

Here, we quantified the functional response of net ecosystem fluxes of oxygen and community structure to temperature, over a broad thermal gradient and in the context of metabolic scaling theory. Oxygen fluxes indicate the productivity of an ecosystem and are directly proportional to carbon fluxes and the potential of an ecosystem to be a carbon source or sink (López-Urrutia *et al.*, 2006). We found that increasing ecosystem average temperatures increased NPP and ER at the community scale, and these effects varied with trophic structure of the local community. The exponential increase in NPP and ER with warming was greatest for communities without consumers (algae only), and least pronounced in communities with grazers and top predators. These results suggest that models relying on functional responses of net ecosystem oxygen or carbon fluxes to temperature might be more accurate when they can include coarse aspects of trophic structure such as the presence of grazers and predators. Further, these results suggest that simplification of trophic structure with environmental change (Estes *et al.*, 2011) could increase the responses of net ecosystem fluxes to temperature changes.

The temperature gradient also affected key aspects of community structure across independent ecosystems, and the response of phytoplankton biomass to temperature was much greater than the effect of temperature on net ecosystem fluxes. These community-level shifts associated with temperature are consistent with what other experimental studies have reported over a much smaller range of temperatures (Beisner *et al.*, 1996; Hansson *et al.*, 2012; Shurin *et al.*, 2012). However, the large community changes over a thermal gradient did not directly predict the effects of temperature on local ecosystem fluxes based on comparisons across ecosystems and metabolic scaling models –variation in fluxes was several orders of magnitude less over the thermal gradient than the variance in phytoplankton biomass. This difference is likely in part attributable to changes in per capita phytoplankton productivity associated with species composition shifts and temperature, and could also be attributed to shifts in phytoplankton abundance relative to benthic algae (which we did not quantify).

We had hypothesized that ecosystem level NPP and ER would be predicted directly by the temperature dependences of photosynthesis and respiration, and insensitive to trophic structure, as they appear to be in many macro-ecological scale analyses (López-Urrutia *et al.*, 2006). However, we found that the temperature dependences varied with trophic structure such that only the grazer-only food chain ecosystems were consistent with temperature dependence of the underlying metabolic processes. Deviations from these expected activation energies could be explained by temperature driven shifts in total biomass. Ecosystem NPP reflected both changes in per capita photosynthesis as well as large changes in mass corrected metabolic biomass (Yvon-Durocher & Allen, 2012), which we were not able to fully characterize in this experiment. Though we did not observe notable amounts of accumulated benthic algae in our tanks, even small amounts could have contributed to total ecosystem fluxes and led to covariation in total biomass with temperature. If the ratio of phytoplankton to benthic algae was temperature-dependent (Dossena *et al.*, 2012), our primary producer biomass estimates may have increasingly under-represented total algal biomass at higher temperatures. In our experiment, we would expect any contribution of benthic algae to NPP to increase over time, and be strongest in weeks 8 and 9 after having had time to accumulate biomass. However, there is no apparent shift in the slopes of NPP *vs* temperature as time progresses (Fig. 2A), suggesting that accumulated benthic algal biomass did not confound our estimate of NPP over the thermal gradient. Nonetheless, to be conservative, we did not present mass-normalized NPP estimates because we could not normalize to any benthic algal metabolic biomass. Covariation between biomass and temperature is common across geographic variation in temperature (Michaletz *et al.*, 2014; Padfield *et al.*, 2017) and therefore present in other estimates of NPP across broad spatial scales when biomass cannot be estimated well.

Another possible reason for the deviation between observed and expected effects of temperature on net oxygen production rates is that resource availability to phytoplankton may have co-varied with temperature such that warmer tanks were less resource-limited. Some cyanobacteria species that increased in our tanks can fix atmospheric nitrogen, but are competitively inferior under conditions of high ambient nitrogen (Hecky & Kilham, 1988). Nitrogen fixation requires the enzyme nitrogenase, which has a biphasic temperature dependence (*E_a_* = −2.18 eV below 22°C and *E_a_* = −0.65 eV above 22°C (Ceuterick *et al*., 1978)). If metabolically active, we speculate that these species may have supplied additional bioavailable N to experimental systems at the warmer end of the thermal gradient (Anderson-Teixeira *et al.*, 2008; Welter *et al.*, 2014). Nutrient limitation may have been eased at higher temperatures by yet another mechanism: all food chains would have included microbial assemblages that may have been recycling nutrients faster at higher temperatures, with respiration-limited metabolic rates (López-Urrutia & Morán, 2007; Beveridge & Humphries, 2010). Fully understanding the effects of temperature on communities and their functions will require including microbial and benthic functional groups.

We observed no sign of ecosystem collapse with warming. Changes in community structure and the increase in trophic control along the temperature gradient appear to be exponential and monotonic (Eqn 1b, Fig. 2), suggesting that observations made from only two temperatures as is typical of many community-level warming experiments may extend to broader thermal gradients using nonlinear (exponential) models (Fig. 2D, Fig. 3). In this pattern, there is little evidence of abrupt transitions that might be expected if thermal stress responses by individual phenotypes emerged at the ecosystem scale. While individual species may experience thermal stress and decline in performance at high temperatures, in our systems these effects were functionally compensated for by other species and increases in per capita performance. The limits to what temperatures could be extrapolated to are not clear from our data, but we would not expect NPP and ER to continue to increase as observed beyond 35C.

Predators, as expected, reduced zooplankton density and body sizes, and caused a clear trophic cascade. Trophic control, and therefore any mitigating effects of predators on biomass change, was weak at low temperatures and increasingly strong at higher temperatures (compare consumer-free control treatment with consumer treatments, Fig. 2D). Over the temperature gradient, community (biomass, abundance) responses were less related to temperature in systems with predators relative to grazer-algae systems. This pattern is consistent with previous findings that systems with two (or even numbers) of trophic levels tend to be more sensitive to warming than systems with odd numbers, due to cascading effects of predation on primary producers (Hansson *et al.*, 2012; Shurin *et al.*, 2012). In our systems, food webs with longer food chains were more resistant to community change with warming. This result contradicts theories in which dynamically responsive predators can make three-trophic-level systems less stable than shorter food chains (Hastings & Powell, 1991). In our experiment, predators were not dynamically responsive. In this way, they represent mortality for zooplankton that may have varied with temperature effects on per capita predation rates by predators, but not demography. In many systems, predators are subsidized by other habitats and food sources, and their populations are not dynamically coupled to prey; in fact, this decoupling has been shown to be important in thermally stratified systems (Tunney *et al.*, 2014). However, our results cannot be extended directly to systems with local dynamic predator population.

Metabolic scaling theory provides an ecological framework that can produce functional responses that relate environmental temperature changes to ecological changes (Padfield *et al.*, 2017). Still missing is a clear understanding of how metabolic temperature dependence emerges at the community level in simple food webs, where effects of species interactions, phenotypic plasticity, evolution and resource limitation can be strong and dominate signals of environmental change (Padfield *et al.*, 2016; 2017). Despite recent advances incorporating light limitation, evolutionary change, trophic interactions and other factors into the metabolic scaling theory framework, empirical tests at the community level such as ours can further shed light on whether there are reliable functional responses of community-scale processes to temperature change. Our study suggests that for oxygen fluxes, metabolic temperature dependence functions associated with MST might be used to model changes with temperature across systems. Changes in species composition and community structure occur within the context set by temperature constraints on energy fluxes via fundamental metabolic processes(Bruno *et al.*, 2015; Padfield *et al.*, 2016; 2017). To extend our findings to a conjecture about implications for climate change, we suggest that conservation actions that maintain predators and top down control may also promote an ecosystem that changes less with temperature than a system with a large abundance of grazers. Taken together, these results suggest our efforts to predict community change with warming may benefit from the general metabolic scaling theory framework to understand even local-scale effects of temperature change at the community level.

## Acknowledgments

We thank W. Cheung and F. Ratcliffe for sampling assistance and D. Song for assistance during zooplankton identification. UBC Mobility Funds to JG, NSERC Discovery Grant to MO.

